# Cognitive effects of early life exposure to PCBs: Sex-specific behavioral, hormonal and neuromolecular mechanisms involving the brain dopamine system

**DOI:** 10.1101/2024.09.13.612971

**Authors:** Emily N. Hilz, Cameron Schnurer, Swati Bhamidipati, Jahnabi Deka, Lindsay M. Thompson, Andrea C. Gore

## Abstract

Endocrine-disrupting chemicals (EDCs) are environmental toxicants that disrupt hormonal and neurodevelopmental processes. Among these chemicals, polychlorinated biphenyls (PCBs) are particularly concerning due to their resistance to biodegradation and tendency to bioaccumulate. PCBs affect neurodevelopmental function and disrupt the brain’s dopamine (DA) system, which is crucial for attentional, affective, and reward processing. These disruptions may contribute to the rising prevalence of DA-mediated neuropsychiatric disorders such as ADHD, depression, and substance use disorders. Notably, these behaviors are sexually dimorphic, in part due to differences in sex hormones and their receptors, which are targets of estrogenic PCBs. Therefore, this study determined effects of early life PCB exposure on behaviors and neurochemistry related to potential disruption of dopaminergic signaling. Male and female Sprague Dawley rats were exposed to PCBs or vehicle perinatally and then underwent a series of behavioral tests, including the sucrose preference test to measure affect, conditioned orienting to assess incentive-motivational phenotype, and attentional set-shifting to evaluate cognitive flexibility and response latency. Following these tests, rats were euthanized, and we measured serum estradiol (E2), midbrain DA cells, and gene expression in the midbrain. Female rats exposed perinatally to A1221 exhibited decreased sucrose preference, and both male and female A1221 rats had reduced response latency in the attentional set-shifting task compared to vehicle counterparts. Conditioned orienting, serum estradiol (E2), and midbrain DA cell numbers were not affected in either sex; however, A1221-exposed male rats displayed higher expression of estrogen receptor alpha (*Esr1*) in the midbrain and non-significant effects on other DA-signaling genes. Additionally, E2 uniquely predicted behavioral outcomes and DAergic cell numbers in A1221-exposed female rats, whereas DA signaling genes were predictive of behavioral outcomes in males. These data highlight sex-specific effects of A1221 on neuromolecular and behavioral phenotypes.

## Introduction

Environmental endocrine-disrupting chemicals (EDCs) are linked to a range of adverse health outcomes in humans and wildlife (Darbre, 2019; Gore et al., 2015; Heindel, 2019; Streifer & Gore, 2021). The developing brain is an especially sensitive target of EDCs (Gore et al., 2015), due to their influences on neurodevelopmental processes that are sculpted by steroid hormones (McCarthy, 2020). Among EDCs, perinatal exposures to polychlorinated biphenyls (PCBs) affect the developing brain’s neuroendocrine systems, with associated behavioral, protein, and gene expression changes observed in the brain of adults (Gillette et al., 2017; Gore et al., 2022; Krishnan et al., 2018). Although banned in the 1970s, PCBs remain environmentally relevant due to resistance to biodegradation and accumulation up the food chain (Borja et al., 2005; Burreau et al., 2004); humans and wildlife throughout the world have detectable PCB body burdens in present times (van den Berg et al., 2017). Epidemiological data suggests PCBs increase the risk of attention-deficit/hyperactivity disorder (ADHD) and associated cognitive disruptions in children and adults (Behforooz et al., 2017; Eubig et al., 2010; Lenters et al., 2019). Despite increasing knowledge about PCB effects on the neuroendocrine system, there are substantial gaps in knowledge on other behaviors and brain systems, including those involved in disorders of attention and hyperactivity related to ADHD, reward-seeking behaviors, and affect.

In the United States, the diagnosis of ADHD has steadily increased in adults and children since the 1990s (Chung et al., 2019; Fairman et al., 2020; Sclar, Robison, Bowen, et al., 2012; Sclar, Robison, Castillo, et al., 2012). The presentation of ADHD is often co-morbid with psychiatric conditions such as mood and substance use disorders, among others (Biederman et al., 2008; Capusan et al., 2019; Chen et al., 2018; Mochrie et al., 2020). The reason for this comorbidity is not well understood but may reflect at least in part shared neurological circuitry that, when disrupted, results in broad changes to behavioral and cognitive function and the presentation of comorbid psychiatric disorders.

Disruption of the brain dopamine (DA) system is a likely candidate to which ADHD and comorbid psychiatric conditions can be attributed: DA synthesis is disrupted in ADHD, higher availability of the DA transporter is observed in ADHD, and the first-choice drugs in treatment for ADHD are DAergic psychostimulants (Faraone, 2018; Krause, 2008; Volkow et al., 2012). DA is an important component of the brain reward system, and it is near dogma that brain DA contributes not only to the hedonic (i.e., pleasurable) aspects of rewarding stimuli, but also and perhaps more importantly to learning and motivation around rewarding experiences (Alikaya et al., 2018; Robinson et al., 2005; Schultz, 2013; Volkow et al., 2009). Anhedonia, or reduced ability to experience pleasure from rewarding stimuli, is a common factor in ADHD, mood disorders (e.g., major depression disorder), and substance use disorder (Der-Avakian & Markou, 2012; Sternat & Katzman, 2016). DA contributes to multiple endophenotypes present in comorbid ADHD including impulsivity, risky decision making, and “bottom-up” executive control wherein self-regulatory processes are heavily informed by subcortical structures (Klein et al., 2019; Volkow et al., 2017).

The DA system develops in early life under the control of estrogens (Ivanova & Beyer, 2003; Kipp et al., 2006; Varshney et al., 2017), making it a likely candidate for estrogenic EDC-mediated disruption which may partially explain the increasing prevalence in ADHD and comorbid psychiatric conditions. Therefore, in the present experiment, we explored the role of exposure to PCBs during sensitive neurodevelopmental windows on hormone and DA-mediated behavioral and physiological endpoints.

## Methods

All procedures conducted on experimental subjects were approved by the Institutional Animal Care and Use Committee at The University of Texas at Austin in accordance with NIH guidelines.

### Subjects

Sexually naive male and female Sprague-Dawley rats (n = 20 / sex) used for breeding were purchased at ∼60 days of age from Envigo (Envigo, Indianapolis, IN, USA). Rats were given 1 week to acclimate to the colony room and were housed under 12:12 hr dark:light conditions with lights off at 11:00 AM, the rats had *ad libitum* access to water and a low phytoestrogen diet (Teklad 2019: Envigo, Indianapolis, IN, USA) unless otherwise noted.

On the day of proestrus, each female was checked for sexual receptivity and then paired overnight with one of the males. The following morning dams were vaginally lavaged; if sperm was present the dams were singly housed, and this was considered embryonic day 0 (E0). From E8 – E18, dams received ‘Nilla wafer cookies treated with either vehicle (Veh; 3% DMSO in sesame oil, Sigma Aldrich, Cat. # D4540 and S3547, respectively) or the PCB mixture Aroclor 1221 (A1221, 1 mg/kg, AccuStandard, Cat # C-221N-50MG), a dose and treatment selected to represent human exposures and because it is a weakly estrogenic EDC in use in the lab for years (Dickerson et al., 2011; Gillette et al., 2022; Reilly et al., 2015). Treatments occurred each morning 2 hours prior to lights out at 1100. Treatments ceased from E19 until birth. On postnatal day 1 (P1), the day after birth, pups (F1) were culled to 6 males and 6 females based on median anogenital index (AGI). Treatments resumed from P1 – P21 through continued feeding of the wafer to the dams. Pups were weaned at P21, after which treatment was discontinued, and littermates were housed in same-sex dyads with weekly handling until behaviors occurred in adulthood (P60). Two F1 male and female pups from each litter were used for behavioral testing (n = 40 / sex).

### Apparati

Sucrose preference testing was conducted in standard polystyrene housing cages measuring 21.5 cm W, 43.5 cm L, and 20 cm H. Steel cage toppers were attached to each cage; two polycarbonate plastic bottles were inserted into the cage topper on either the left or right side, and low phytoestrogen diet was provided *ad libitum* in the center. The bottles contained either tap water or 0.08% sucrose solution.

Pavlovian light-food conditioning and attentional set-shifting were performed in standard conditioning chambers with dimensions of 30.5 cm W, 25.5 cm L, and 30.5 cm H (Coulbourn Instruments, Whitehall, PA). The chambers featured clear acrylic walls at the front and back, steel-rod flooring, and aluminum sides and ceiling. Food pellets (45 mg TestDiet, Richmond, IN) acted as an unconditioned stimulus (US) and were dispensed into a food cup on the right wall via an external magazine. Entries into the food cup were measured via infrared beam at the cup’s opening. A 2-watt bulb positioned 20 cm above the food cup served as the conditioned stimulus (CS) when illuminated. Five open ports were located on the left side of the chamber; for Pavlovian conditioning, the ports were inactive and covered. For attentional set-shifting the outer left, right, and center ports were blocked and were not used during the procedure. Nose-pokes into the ports were measured via infrared beam at the opening of each port. The chambers were housed in sound-and light-attenuating enclosures (Coulbourn Instruments, Whitehall, PA). Digital video cameras (KT&C USA, Fairfield, NJ), mounted inside the enclosures but outside the chambers, were used to record orienting activity during Pavlovian food-light conditioning.

### Behavioral Procedures

#### Sucrose preference testing

Prior to the procedure, sucrose and tap water bottles were weighed. Rats were placed individually into a clean standard housing cage, and then both sucrose and tap water bottles were inserted into the cage topper with the location of the bottles (left or right) semi randomized. The procedure began at 08:00 AM and lasted 24 hours; at the 12-hour timepoint, the location of the sucrose and tap bottle were switched to account for potential side biases. The following day, rats were removed from the test cages at 08:00 AM and bottles were weighed. The difference in weight (g) from beginning to end of the procedure was used to generate sucrose preference scores by dividing the amount of sucrose solution consumed by the total liquid (sucrose and tap) consumed over the 24-hour period.

#### Pavlovian Conditioning

Prior to Pavlovian conditioning, rats were food restricted for 5 days to reach 90% body weight. Food restriction was maintained throughout the Pavlovian conditioning and attentional set-shifting procedures, and both procedures were conducted under red light beginning 1 hour after lights out (12:00 PM).

On the first day of Pavlovian conditioning, rats underwent training to retrieve food pellets from the foodcup where 30 food unconditioned stimuli (USs) were delivered on a 60-second fixed inter-trial interval (ITI) with no light conditioned stimulus (CS). The following day, rats underwent habituation to the light CS. For the first 8 trials, the light CS was presented for 10 seconds without delivery of the food US; for the second 8 trials, the 10-second light CS preceded delivery of 1 food US. Habituation occurred over ∼35 minutes with a variable ITI of 120 seconds +/-60 seconds. The subsequent conditioning sessions occurred over the following 3 days and consisted of 16 CS-US presentations on the same ITI schedule. Each session video recording was scored by a blind and independent observer. A conditioned orienting response (OR) was defined as vertical rearing of the rat wherein the front paws leave the floor, excluding grooming behavior (Hilz et al., 2021; Hilz, Lewis, et al., 2019; Hilz, Smith, et al., 2019). Because the light CS is diffuse in the conditioning chamber, ORs were counted regardless of if the body was oriented towards or away from the CS. Each CS presentation was divided into 3 5-second (s) intervals: pre-CS: 5 s prior to CS illumination (a baseline measure of OR), CS1: the first 5 s of the CS, and CS2: the second 5 s of the CS. ORs were scored every 1.25 s, allowing for up to 4 OR scores in each CS. Pre-CS OR scores were subtracted from CS1 and CS2 OR scores, providing a final OR score adjusted for any unconditioned ORs occurring prior to the CS illumination.

#### Attentional Set-Shifting

On the first day of attentional set-shifting, rats were trained to nose-poke into one of two open ports on the left side of the conditioning chamber. Each trial was signaled by illumination of the house light; after 3 seconds, one of the two open ports was illuminated with red light and the rat was required to make a nose-poke into the illuminated port within 10-seconds to receive 1 food pellet. This training lasted until the rats made 50 successful nose-pokes within a 30-minute period, which was completed for all rats in one training session. The location of the illuminated port was semi-randomized such that half of the rats were trained beginning with the left port and half of the rats were trained beginning with the right port.

The following day, rats underwent the set-shifting procedure. The procedure was counter-balanced such that half of the rats began with the response discrimination task, and half began with the visual discrimination task. For both tasks, the beginning of a trial was signaled with a 3-second illumination of the house light and subsequently one of the two open ports would illuminate for up to 10 seconds. Trials were presented on a variable 20-second ITI, and ports were illuminated semi-randomly such that neither port illuminated more the 3 times consecutively.

For the spatial discrimination task, rats were required to nose-poke one of the two ports (left or right) regardless of whether the port was illuminated. Half of the rats were required to nose-poke the left port, and half of the rats were required to nose-poke the right port. A nose-poke to the correct port resulted in delivery of 1 food pellet and darkening of the port and house light. Nose-poking to the incorrect port or failing to nose-poke to either port within 10-seconds (counted as omissions) resulted in darkening of the house light and port, and no food pellet delivery. The test lasted for up to 2 hours and completed after the rat met a criterion of 10 correct responses in a row. Most rats completed the task within one session, but those that did not were required to undergo the task again the following day until criterion was met. After reaching criterion, rats that began testing with the spatial discrimination task underwent the visual discrimination task on the following day.

For the visual discrimination task, rats were required to nose-poke the illuminated port regardless of its location in space (left or right). Otherwise, the experimental parameters were identical to the response discrimination task. After reaching the criterion of 10 correct consecutive responses, rats that began testing with the visual discrimination task underwent the spatial discrimination task on the following day.

### Euthanasia and collection of biological specimens

Rats were allowed to eat *ad libitum* and regain body weight for one week prior to euthanasia. At ∼P80 rats were euthanized by either rapid decapitation or perfusion; females were euthanized during proestrus based on cytological examination of cells collected via vaginal lavage. For the subset of rats that were rapidly decapitated (n = 20 / sex), brains were snap-frozen in isopentane, and trunk blood was collected, allowed to clot, and centrifuged to separate out serum. Both brains and serum were stored at -80 °C until use. For the subset of rats that were perfused (n = 20 / sex), the rats were overdosed with an intraperitoneal injection of ketamine (100 mg/kg) and xylazine (40 mg/kg). Once anesthetized, cardiac blood was collected, allowed to clot, centrifuged to separate out serum, and stored at -80 °C. The rats were then perfused for 1 minute with 1% paraformaldehyde (PFA) in phosphate buffered saline (PBS), and then for 8 minutes with 4% PFA in PBS. Brains were stored for 24 hours at 4 °C in 4% PFA-PBS, and then underwent 3 days of gentle agitation in increasing sucrose solution concentrations (10%, 15%, and 30%) until finally being stored at -20 °C in cryoprotectant.

### Serum estradiol radioimmunoassay

Circulating serum estradiol (E2) from all behaviorally characterized rats was quantified using radioimmunoassay (RIA) according to the manufacturer’s protocol (Beckman Coulter, DSL4800). The RIA was conducted in a single cohort and samples were analyzed in duplicate; the sensitivity of the assay was 2.2 pg/mL with an average intra-assay CV of 5.67%.

### Tissue processing and dopamine immunohistochemistry

Perfused brains (*n* = 20 / sex; 40 total) were sliced coronally into 6 series at 30 µm on a vibrating microtome (Leica VT 1000S). The samples were stored at -20 °C in cryoprotectant until immunohistochemical (IHC) processing. One series containing the substantia nigra and ventral tegmental areas of the midbrain was processed using IHC for tyrosine hydroxylase (TH), the rate-limiting enzyme in catecholamine biosynthesis that is commonly used to identify DAergic cells in DA-producing regions of the brain. Briefly, tissues were washed in PBS followed by 0.3% H_2_O_2_ in PBS; tissues were gently agitated for 1 hour in blocking buffer (6% normal goat serum in 0.3% PBS-Triton) and then underwent primary incubation with TH antibody (1:2500; 22941, ImmunoStar, Hudson, WI) in blocking buffer (3% NGS in 0.15% PBS-T), gently agitated overnight at 4 °C. Tissues were washed, incubated in secondary biotinylated goat anti-mouse IgG (1:250; Vector Laboratories) for 1 hour, and then VECTASTAIN Elite ABC-HRP Kit (Vector) was used and tissue was stained with 3’3 diaminobenzidine (Vector). The tissue was mounted onto Superfrost Plus slides (Thermo Fisher Scientific, Carlsbad, CA, USA), allowed to dry overnight, dipped in a series of increasing EtOH concentrations followed by xylene, and cover-slipped using DPX mounting media (Fisher).

Samples were imaged at 20x magnification using an Olympus BX61 bright field microscope. Each sample was imaged bilaterally; for the SN, images were captured at -4.80 mm, -4.92 mm, and -5.04 mm from bregma (Paxinos & Watson, 1986) and two images were taken at each level. For the VTA, images were captured at -5.04 mm from bregma and one image was taken at this level. The number of TH+ cells for each image were quantified by an observer blinded to the experimental condition.

### Tissue processing and qPCR

Fresh frozen brains (*n* = 20 / sex; 40 total) were sliced coronally on a NX50 cryotstat (Fisher) at 400 µm. SN and VTA regions were immediately punched bilaterally at 0.75 mm using a Palkovits punch (Stoelting, Wood Dale, IL, USA). The SN and VTA punches were combined in a chilled microfuge tube and stored at -80 °C. RNA was extracted using AllPrep DNA/RNA/miRNA Universal Kit (80224; Qiagen, Germantown, MD) according to the manufacturer’s protocol with added DNase. The concentration of RNA was confirmed by a nanodrop. RNA was concentrated via vacuum centrifugation, resuspended in nuclease free water, and diluted to 20 ng/µl for cDNA conversion (High Capacity cDNA Reverse Transcription Kit, Thermo Fisher Scientific). Real-time PCR was performed with TaqMan Gene Expression Master Mix (Thermo Fisher Scientific) using ViiA7 and QuantStudio 7 Real-time PCR Systems (Applied Biosystems, Carlsbad, CA, USA) at 50 °C (2 min), 95 °C (10 min), and 42 cycles of 95 °C (15 s) and 60 °C (1 min). Primers and probes were purchased from Thermo Fisher Scientific for estrogen receptor alpha (*Esr1*), estrogen receptor beta (*Esr2*), dopamine receptor 1 (*Drd1*), dopamine receptor 2 (*Drd2*), dopamine transport gene *Slc6a3*, and tyrosine hydroxylase coding gene *Th*. The primers and probes were duplexed to the housekeeping gene *Gapdh*, and relative gene expression was determined using the comparative Ct method wherein samples were normalized to *Gapdh* and calibrated to the median delta-cycle threshold of Veh controls (ΔΔCt). Fold changes in gene expression were calculated using the formula: 2^-(ΔΔCt).

### Statistical Analysis

All statistical analyses were performed in R studio version 4.3.1 (R Core Team, 2023). Unless otherwise noted, outcome variables were analyzed using within-subjects 2×2 ANOVAs for repeated measures data or between-subjects 2×2 ANOVAs for non-repeated measures data using the common factor “Treatment” (i.e., Veh or A1221). ANOVAs were applied separately within each sex. Normality and homogeneity of variance in the data were first confirmed using Shapiro-Wilk test of normality and Bartlett’s test of homogeneity of variance; data that violated assumptions were transformed using optimized Box-Cox power transformations. Appropriate *post hoc* analyses were applied to significant main or interaction effects: Bonferroni correction for repeated measures data, otherwise Tukey HSD; a measure of effect size, partial eta squared (n ^2^), was provided for significant comparisons.

#### Behavioral analyses

For the sucrose preference test, the factor “Solution” indicated the amount of tap water or sucrose solution consumed in grams, and “Preference” indicated the percentage of sucrose solution consumed relative to total liquid consumed. For Pavlovian conditioning, the factor “Block” indicated acquisition of ORs over time (i.e., blocks of 8 trials, 4 days) and the first block, which represented unpaired CS-US presentations, was excluded from analysis. For the attentional set-shifting task, the factor “Condition” indicated the initial attentional “set” building task (i.e., spatial or visual discrimination task, counterbalanced) and the subsequent “shift” to the new requirement task (i.e., spatial or visual discrimination task, counterbalanced). “Counterbalance” indicated the order in which rats underwent the tasks (i.e., spatial-to-visual or visual-to-spatial), and the response variables analyzed were nose pokes into the correct or incorrect port, and latency to respond.

#### Biological measures

E2 concentration, the number of TH+ cells quantified in the SN and VTA, and expression of target genes were analyzed with factorial ANOVAs using the factors “E2” (i.e., mean serum E2 concentration from sample duplicates), “DA” (i.e., mean number of TH+ cells from samples quantified), or factors relative to the target gene (i.e., mean of fold gene expression for *Esr1, Esr2, Drd1, Drd2, Slc6a3,* and *Th*).

#### Regression analyses

Regression models were generated for each sex separately using the ‘lm’ function from the ‘stats’ package in R. For each, the dependent variable matched factors described in *behavioral analyses*; E2 (all rats, *n* = 20 / treatment / sex; 80 total), DA cells in either SN or VTA (from the subset of perfused rats, *n* = 10 / treatment / sex; 40 total), and each of the target genes from SN and VTA combined (from the subset of rats that were rapidly decapitated, *n* = 10 / treatment / sex; 40 total) acted as independent variables, and treatment group was included as an interaction term in each model. Data were converted to z-scores prior to analysis. The overall model was considered significant if *p* < 0.05; beta coefficient (β), standard error (SE), t-value, and p-value were reported for significant main effects and/or interaction terms.

## Results

### Behavioral outcomes

#### Sucrose Preference Test

All rats consumed more sucrose solution than tap water (Figure 1A), indicated by a main effect of Solution in both female (*F*(1,38) = 182.88, *p* < 0.001, n ^2^ = 0.83; *post hoc* adjusted *p* < 0.001) and male rats (*F*(1,38) = 168.69, *p* < 0.001, n ^2^ = 0.82; *post hoc* adjusted *p* < 0.001). In males, there was no effect of Treatment nor interaction between Treatment and Solution (*p* > 0.1 for both). In females, a non-significant main effect of Treatment (*F*(1,38) = 3.12, *p* = 0.08, n ^2^ = 0.07) and a non-significant interaction between Treatment and Solution (*F*(1,38) = 3.36, *p* = 0.07, n ^2^ = 0.08) were detected; although these did not meet conventional levels of significance and the effect size was small, *post hoc* analysis indicated marginally higher consumption of tap water in the A1221 females compared to Veh controls (*p* = 0.07). Preference for sucrose solution was lower in female rats exposed to A1221 compared to Veh (*F*(1,38) = 4.16, *p* = 0.05, n ^2^ = 0.10; *post hoc* adjusted *p* = 0.05; Figure 1B), but was not different between male treatment groups (*p* > 0.1; Figure 1B).

**Figure 1.**
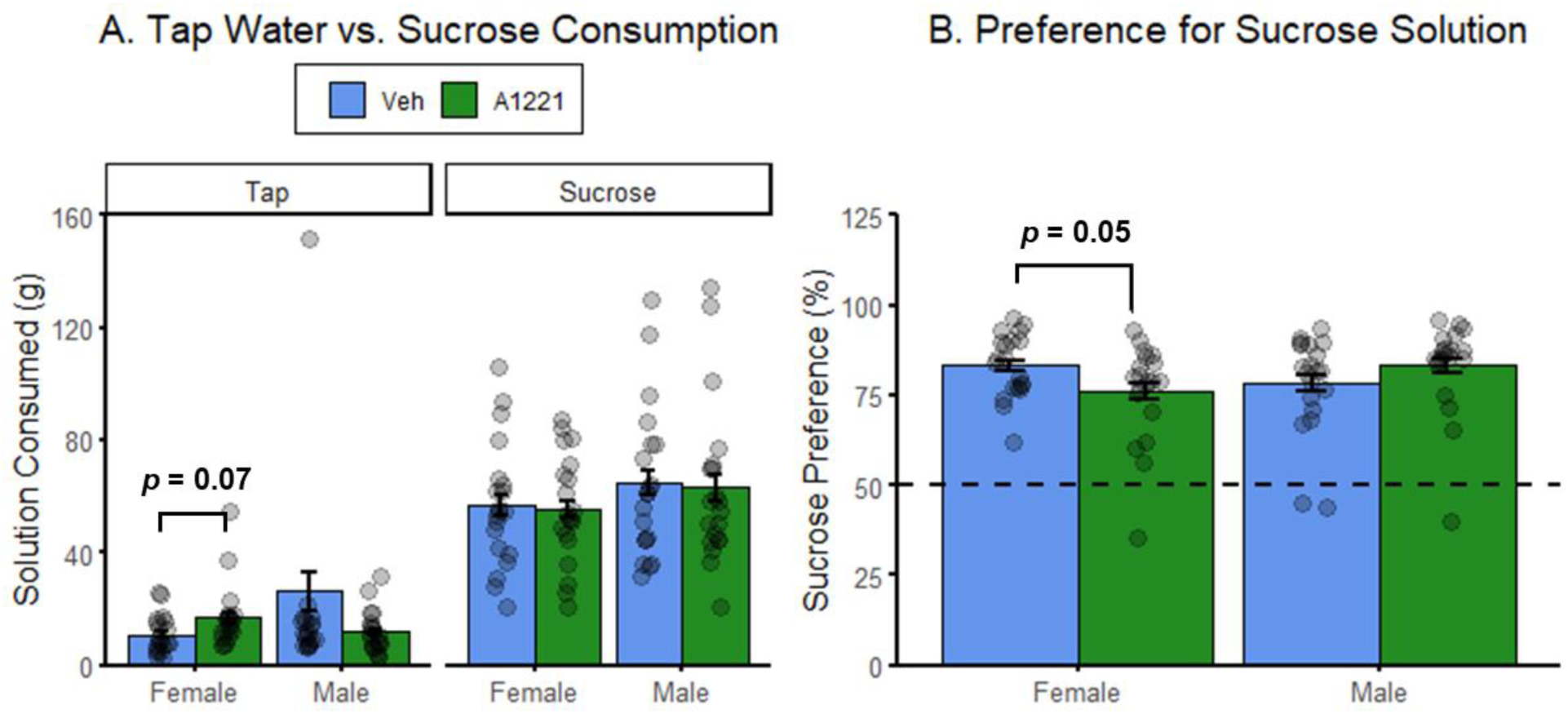
Sucrose preference test results. Points indicate individual scores, N = 20/sex/treatment. **A.** Mean tap water or sucrose solution consumed in grams (g) +/- SEM. Female rats with A1221 exposure drank a non-significantly higher amount of tap water (left panel) compared to Veh controls (*p* = 0.07). There was no difference in sucrose solution consumption between treatment groups in either sex. **B**. Mean percent (%) preference for sucrose solution +/- SEM. The dashed line indicates the 50% preference point, above which rats exhibit increased preference for the sucrose solution. Female rats with A1221 exposure had a significantly lower preference for sucrose solution compared to Veh controls (*p* = 0.05). Male sucrose preference was not different between treatment groups.

#### Conditioned Orienting

We previously found that OR behavior was consistently higher in CS1 and foodcup (FC) behavior was higher in CS2 (Hilz et al., 2021; Hilz, Lewis, et al., 2019; Hilz, Smith, et al., 2019; Olshavsky et al., 2014). The current results mirrored our previous findings: in both females and males, OR behavior was higher in CS1 and FC behavior was higher in CS2 (Figure 2A,B). In females, a non-significant main effect of CS (*F*(1,501) = 3.32, *p* = 0.07, n ^2^ = 0.01) and an interaction between CS and Block (*F*(6,501) = 4.79, *p* < 0.001, n ^2^ = 0.06) indicated that ORs were marginally higher overall in CS1 compared to CS2 (*p* = 0.07) and were significantly higher in CS1 by the end of conditioning (*p* = 0.006). In males the data followed a similar but more robust pattern: a main effect of CS (*F*(1,503) = 37.73, *p* < 0.001, n_p_^2^ = 0.07) and an interaction between CS and Block (*F*(1,503) = 11.98, *p* < 0.001, n_p_^2^ = 0.13) indicated that ORs were higher overall in CS1 (*p* < 0.001) and by end of conditioning (*p* < 0.001). For FC behavior, females had a main effect of CS (*F*(1,507) = 162.79, *p* <0.001, n ^2^ = 0.24) and an interaction between CS and Block (*F*(6,507) = 9.78, *p* < 0.001, n ^2^ = 0.10), indicating higher FC behavior in CS2 both overall and at the end of conditioning (*p* < 0.001 for both). Males also had a main effect of CS (*F*(1,503) = 37.73, *p* < 0.001, n ^2^ = 0.07) and an interaction between CS and Block (*F*(1,503) = 11.98, *p* < 0.001, n ^2^ = 0.13), similarly indicating higher FC behavior in CS2 both overall and at the end of conditioning (*p* < 0.001 for both).

**Figure 2.**
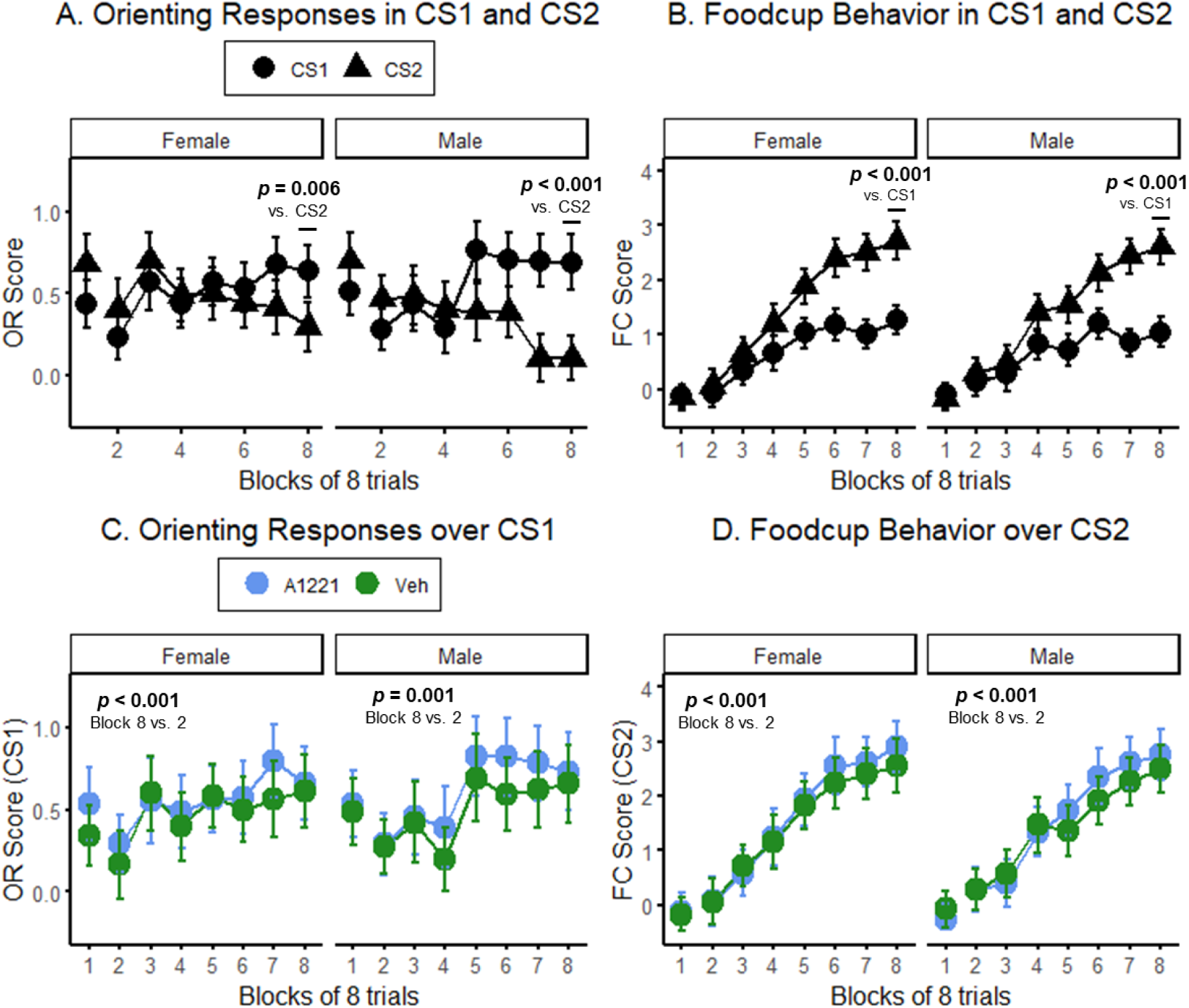
Orienting response (OR) and foodcup (FC) scores over Pavlovian conditioning, N = 20/sex/treatment. **A.** Mean OR scores over conditioning in CS1 and CS2 +/- SEM. In both female (left panel) and male rats (right panel), OR scores were higher at the end of conditioning (block 8) in CS1 compared to CS2 (*p* = 0.006 and *p* < 0.001, respectively). **B.** Mean FC scores over conditioning in CS1 and CS2 +/- SEM. In both sexes, FC scores were higher at the end of conditioning (block 8) in CS2 compared to CS1 (*p* < 0.001 for both). **C.** Mean CS1 OR scores over conditioning +/- SEM. In both sexes, OR scores were higher in block 8, the end of conditioning, compared to the start of conditioning at block 2 (*p* < 0.001 and *p* = 0.001, respectively). There were no effects of treatment group. **D.** Mean CS2 FC scores over conditioning +/- SEM. In both sexes, FC scores were higher at the end of conditioning compared to the beginning (*p* < 0.001 for both). There was no effect of treatment group.

Both female and male rats significantly acquired ORs and FC behavior over the course of Pavlovian conditioning (OR: *F*(6,226) = 6.36, *p* < 0.001, n ^2^ = 0.14 and *F*(6,226) = 9.90, *p* < 0.001, n ^2^ = 0.21, respectively; FC: *F*(6,228) = 51.57, *p* < 0.001, n ^2^ = 0.57 and *F*(6,228) = 36.04, *p* < 0.001, n ^2^ = 0.49; Figure 2C,D). *Post hoc* analyses indicated that female and male rats had higher OR and FC scores at the end of conditioning compared to the beginning (OR: *p* < 0.001 and p = 0.001, respectively; FC: *p* < 0.001 for both). There was no effect of Treatment on the acquisition of ORs nor on FC behavior in either sex (*p* > 0.1 for all comparisons), nor on OR or FC level at the end of conditioning (*p* > 0.1 for both sexes).

#### Attentional set-shifting

The total number of trials required to reach criterion was significantly higher after the shift in response requirements in both female (*F*(1,36) = 21.67, *p* < 0.001, n ^2^ = 0.38; *post hoc* adjusted *p* < 0.001; Figure 3A) and male rats (*F*(1,36) = 18.42, *p* < 0.001, n ^2^ = 0.34; *post hoc* adjusted *p* < 0.001; Figure 3A). Neither sex showed an effect of Treatment nor interaction between Treatment and Condition (*p* > 0.1 for all comparisons). In females, the order of tests did not influence overall trials required to reach criterion (*p* > 0.1); however, male rats in the spatial-to-visual test order group required significantly more trials to reach criterion after shifting response requirements than male rats in the visual-to-spatial group (*F*(1,36) = 12.03, *p* = 0.001, n ^2^ = 0.25; *post hoc* adjusted *p* < 0.001). Incorrect responses followed a similar pattern: females made more incorrect responses before reaching criterion after the shift in response requirements (*F*(1,36) = 17.40, *p* < 0.001, n ^2^ = 0.33; *post hoc* adjusted *p* < 0.001; Figure 3B) without any test order effect (*p* > 0.1). Males also made more incorrect responses after shifting response requirements (*F*(1,36) = 16.09, *p* < 0.001, n ^2^ = 0.31; *post hoc* adjusted *p* = 0.001; Figure 3B), and incorrect responses were higher in males in the spatial-to-visual test order group compared to the visual-to-spatial group (*F*(1,36) = 12.73, *p* = 0.001, n ^2^ = 0.27; *post hoc* adjusted *p* < 0.001). There were no effects nor interactions of Treatment on incorrect responses in either sex (*p* > 0.1 for all comparisons).

**Figure 3.**
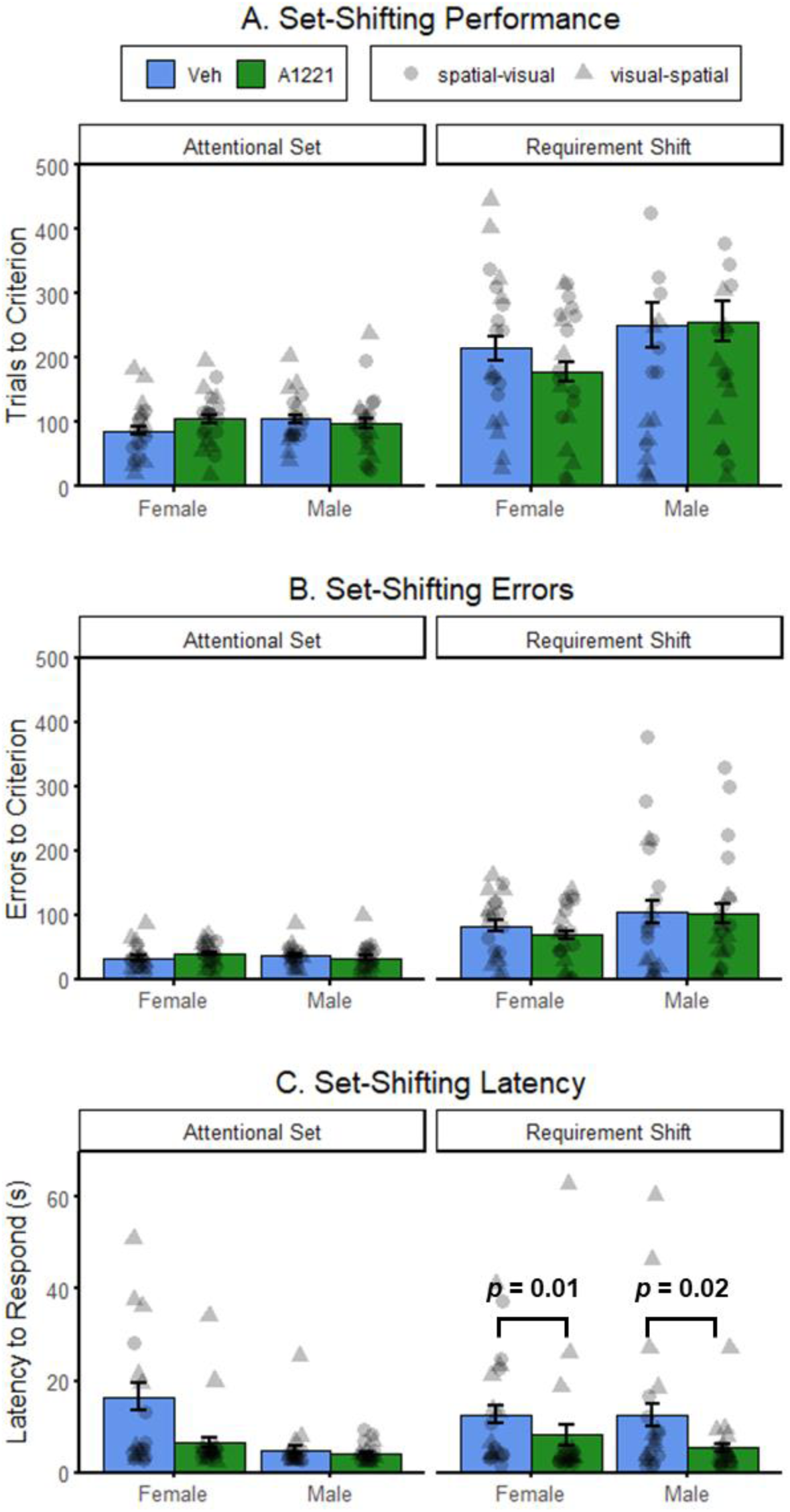
Response measures in the attentional set-shifting task. Points indicate individual scores, shape indicates counterbalanced order of response requirements. N = 20/sex/treatment. **A.** Mean total trials to criterion +/- SEM. In both female and male rats, the total trials required to reach criterion did not differ between treatment groups in the attentional set task (left panel) nor after the shift in response requirements (right panel). **B.** Mean errors before criterion +/- SEM. In both female and male rats, errors emitted prior to reaching criterion did not differ between treatment groups in the attentional set task (left panel) nor after the shift in response requirements (right panel). **C.** Mean latency to respond in seconds (s) +/- SEM. In both female and male rats, A1221 exposure decreased response latency compared to sex-specific Veh controls (*p* = 0.01 and *p* = 0.02, respectively).

An effect of Treatment was observed in the latency to make a response. After the shift in response requirements, latency to respond was lower in both A1221-exposed males (*F*(1,36) = 5.59, *p* = 0.02, n ^2^ = 0.13; *post hoc* adjusted *p* = 0.02; Figure 3C) and A1221-exposed females (*F*(1,36) = 6.80, *p* = 0.01, n ^2^ = 0.16; *post hoc* adjusted *p* = 0.01; Figure 3C) compared to sex-specific Veh controls. The order of test presentation was a significant factor in both females (*F*(1,36) = 6.79, *p* = 0.01, n_p_^2^ = 0.16; *post hoc* adjusted *p* = 0.01) and males (*F*(1,36) = 14.47, *p* < 0.001, n_p_^2^ = 0.29; *post hoc* adjusted *p* < 0.001), such that response latency was higher in the visual-to-spatial test order group for both sexes. The order of tests did not interact with Treatment to affect response latency (*p* > 0.1 for all).

### Physiological measures

#### Serum estradiol

Serum E2 levels did not differ between treatment groups within female (p > 0.01) nor male rats (p > 0.1; Figure 4A).

**Figure 4.**
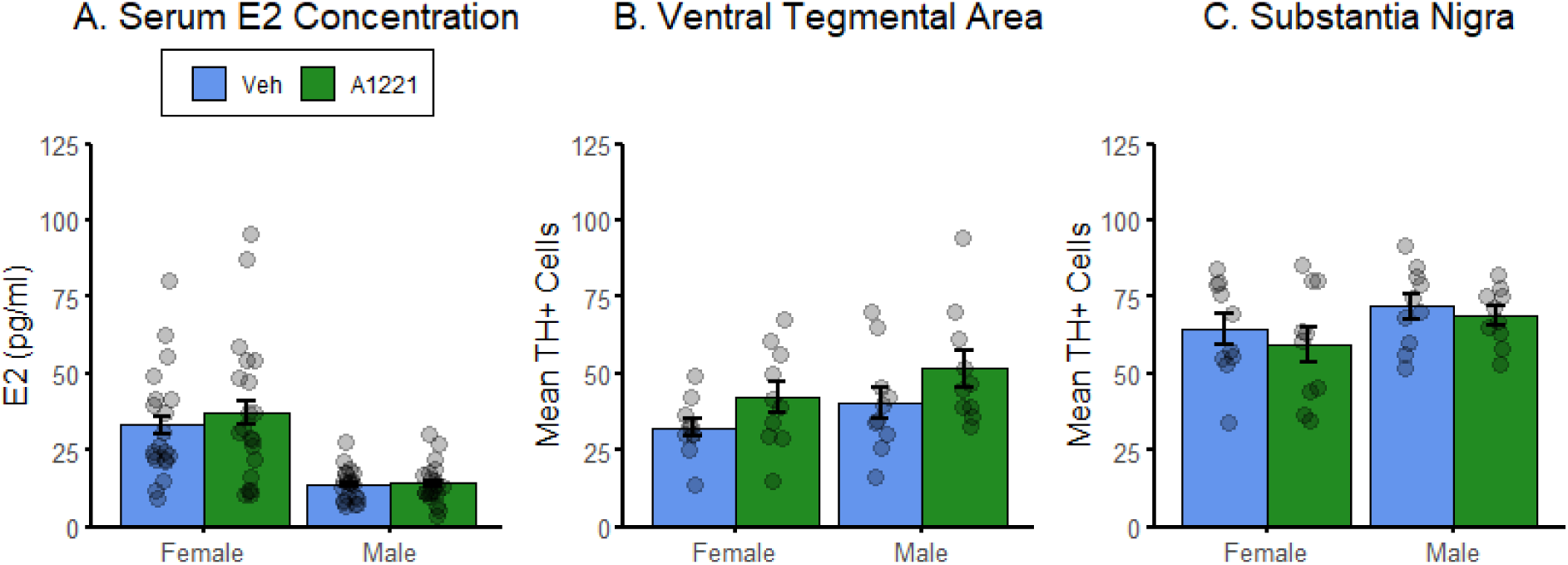
Serum estradiol (E2; N = 20/sex/treatment) and tyrosine hydroxylase positive (TH+; N = 10/sex/treatment) cells in the midbrain Ventral Tegmental Area and Substantia Nigra. Points indicate individual scores. **A.** Mean serum E2 (pg/ml) +/- SEM. In all behaviorally characterized female and male rats (*n* = 20 / sex / treatment), serum E2 did not differ between treatment groups. **B.** Mean TH+ cells in the Ventral Tegmental Area (VTA) +/- SEM. In the subset of rats used for immunohistochemistry (*n* = 10 / sex / treatment), TH+ cell numbers did not differ between treatment groups in the VTA. **C.** Mean TH+ cells in the Substantia Nigra (SN) +/- SEM. In the subset of rats used for immunohistochemistry (*n* = 10 / sex / treatment), TH+ cell numbers did not differ between treatment groups in the SN.

#### Midbrain dopamine cell numbers

In the VTA, the mean number of TH+ cells did not differ between treatment groups in female (*p* = 0.1; Figure 4B) nor male rats (*p* > 0.1; Figure 4B). Similarly, in the SN, the mean number of TH+ cells did not differ between treatment groups in female (*p* > 0.1; Figure 4C) nor male rats (*p* > 0.1; Figure 4C).

### Gene expression analyses

Expression of six genes was measured in the combined VTA and SN of individual rats (Figure 5). *Esr1* was not different between Veh and A1221-exposed females (*p* > 0.1); however, males with perinatal A1221 exposure had significantly higher *Esr1* expression compared to Veh controls (*F*(1,18) = 5.57, *p* = 0.03, n ^2^ = 0.24; *post hoc* adjusted *p* = 0.03; Figure 5A). *Drd1* was not different between Veh and A1221-exposed females (*p* > 0.1); however males had a non-significant effect of Treatment on *Drd1* expression (F(1,18) = 3.87, p = 0.06, n ^2^ = 0.18; Figure 5C). Although this did not meet conventional levels of significance, the effect size was strong and *post hoc* analyses indicated that A1221 males had marginally higher *Drd1* expression compared to Veh controls (*post hoc* adjusted *p* = 0.06). Similarly, DA transporter gene *Slc6a3* was not different between Veh and A1221-exposed females (*p* > 0.1); however, males had a non-significant effect of Treatment on *Slc6a3* expression (*F*(1,18) = 3.24, *p* = 0.08, n ^2^ = 0.15; Figure 5E). Although this did not meet conventional levels of significance, the effect size was strong and *post hoc* analysis indicated marginally lower expression of *Slc6a3* in the A1221 males compared to Veh controls (*post hoc* adjusted *p* = 0.08). *Drd2*, *Esr2*, and the tyrosine hydroxylase encoding gene *Th* were not different between treatment groups in either sex (*p* > 0.1 for all comparisons; Figure 5B,D,F).

**Figure 5.**
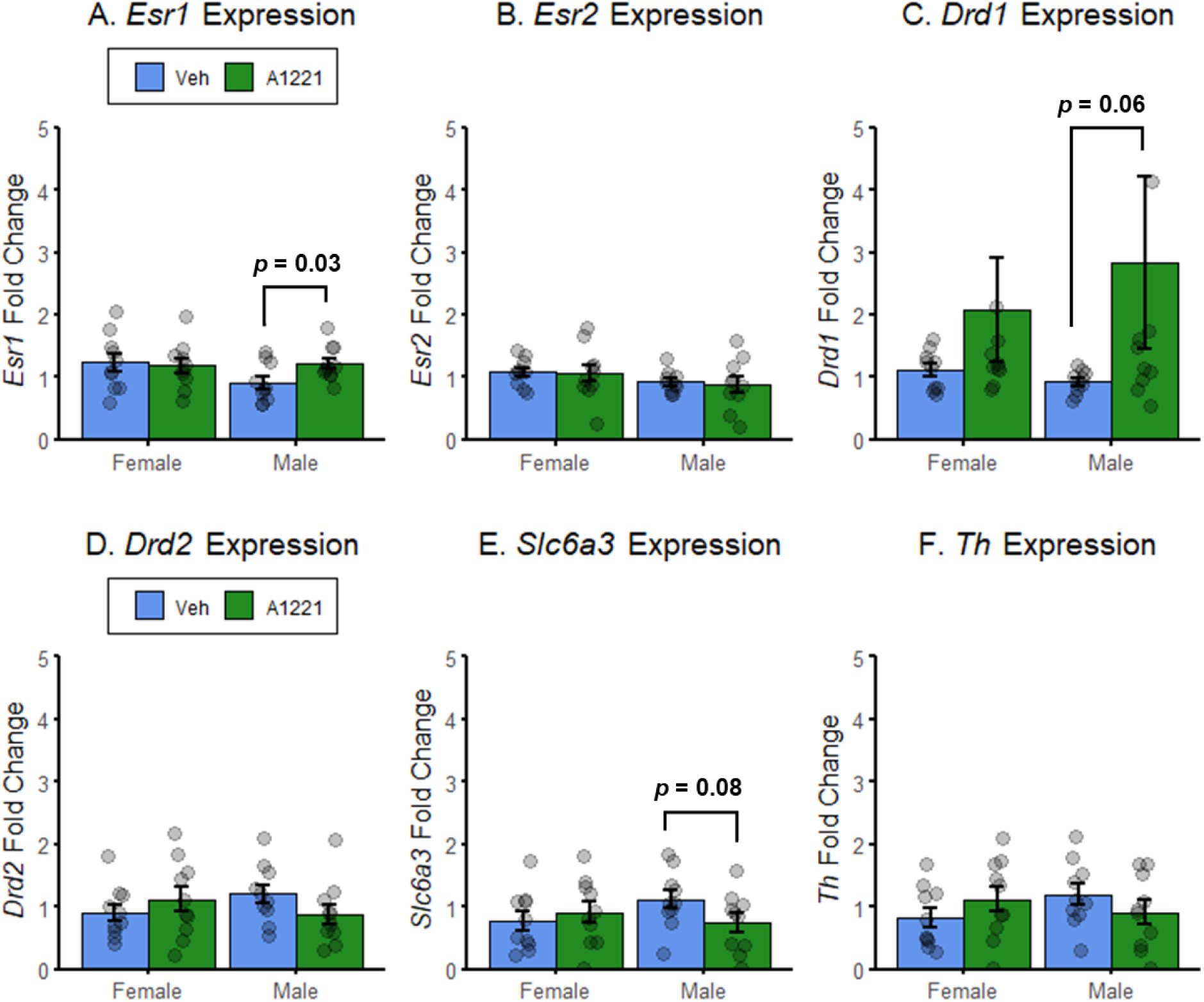
Expression of target genes in the midbrain (substantia nigra and ventral tegmental area combined). Points indicate individual scores. N = 10/sex/treatment. **A.** Mean fold change in estrogen receptor alpha (*Esr1*) +/- SEM. In male rats, A1221 exposure increased *Esr1* expression relative to Veh controls (*p* = 0.03). **B.** Mean fold change in estrogen receptor beta (*Esr2*) +/- SEM. In female and male rats, *Esr2* expression did not differ between treatment groups. **C.** Mean fold change in dopamine receptor 1 (*Drd1*) +/- SEM. In male rats, A1221 exposure non-significantly increased *Drd1* expression relative to Veh controls (*p* = 0.06). **D.** Mean fold change in dopamine receptor 2 (*Drd2*) +/- SEM. In female and male rats, *Drd2* expression did not differ between treatment groups. **E.** Mean fold change in dopamine transport gene *Slc6a3* +/- SEM. In male rats, A1221 exposure non-significantly decreased *Slc6a3* expression relative to Veh controls (*p* = 0.08). **F.** Mean fold change in tyrosine hydroxylase encoding gene *Th* +/- SEM. In female and male rats, *Th* expression did not differ between treatment groups.

### Regression analyses

#### Estradiol and behavior

In females, serum E2 predicted the total number of responses required to reach criterion in the set-shifting task after the response requirement shift differently between treatment groups (Figure 6A). The overall model explained 24.2% of the variation in the data, with an adjusted R^2^ of 17.9% (*F*(3,36) = 3.84, *p* = 0.02). The relationship between E2 and total responses was negative (β = -0.25, STE = 0.11, *t* = -2.36, *p* = 0.02); E2 interacted positively with Treatment and responses (β = 0.55, STE = 0.18, *t* = 3.12, *p* = 0.004), indicating that E2 was a positive predictor of performance in the set-shifting task in Veh controls, but not in A1221-exposed females. A similar model showed a predictive relationship between serum E2 and incorrect responses after the requirement shift in female rats (Figure 6B). The overall model explained 24.6% of the variation in the data, with an adjusted R^2^ of 18.4% (*F*(3,36) = 3.93, *p* = 0.02). The relationship between E2 and incorrect responses was negative (β = -0.23, STE = 0.09, *t* = -2.38, *p* = 0.02); E2 interacted positively with Treatment and responses (β = 0.52, STE = 0.16, *t* = 3.21, *p* = 0.003), indicating that E2 was a positive predictor of performance in the set-shifting task in Veh controls, but not in A1221-exposed females. E2 did not predict behavioral outcomes in male rats (all models *p* > 0.1).

**Figure 6.**
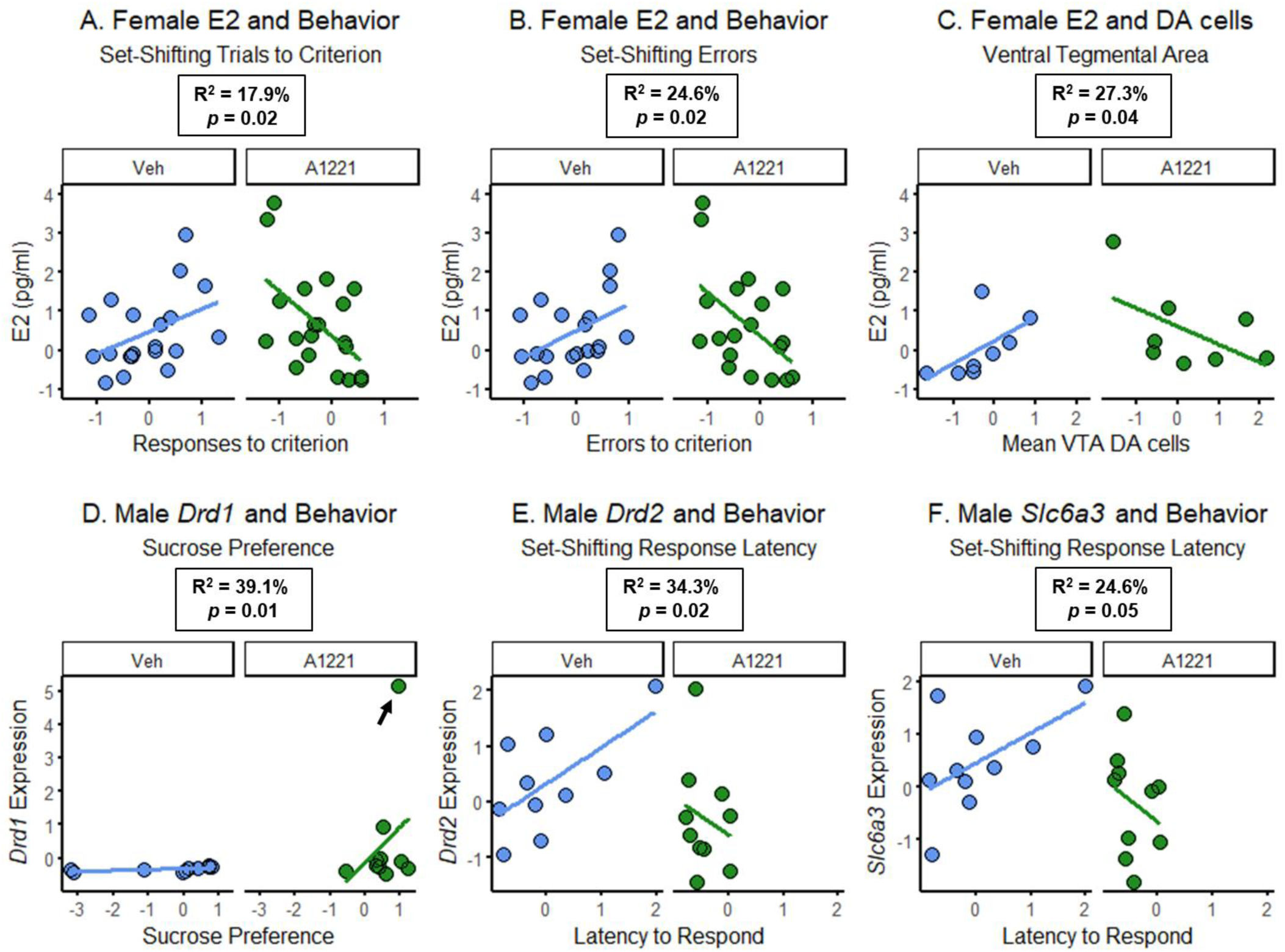
Scatterplots with regression lines showing significant relationships between outcomes variables. Points indicate individual scores, and lines indicate trends within treatment groups; all data are standardized to z-scores. R^2^ and *p*-values are from the omnibus regression analysis are shown. **A.** Serum estradiol (E2; N = 20/ treatment) explained 17.9% (R^2^) of the variation in the trials required to reach criterion after the response requirement shift in female rats (*p* = 0.02). The slope of the regression lines were significantly different between A1221 females and Veh controls. **B.** Serum estradiol (E2) explained 24.6% (R^2^) of the variation in errors before reaching criterion after the response requirement shift in female rats (*p* = 0.02). The slope of the regression lines were significantly different between A1221 females and Veh controls. **C.** In the subset of female rats used for IHC (N = 10 / treatment group), serum estradiol (E2) explained 24.6% (R^2^) of the variation in Ventral Tegmental Area (VTA) dopamine (DA) cells (i.e., TH+ cells; *p* = 0.02). The slope of the regression lines were significantly different between A1221 females and Veh controls. **D.** In the subset of male rats used for PCR (N = 10 / treatment group), expression of dopamine receptor 1 (*Drd1*) from the combined midbrain regions explained 39.1% (R^2^) of the variation in preference for sucrose solution (*p* = 0.01). The slope of the regression lines were significantly different between A1221 males and Veh controls. The arrow indicates a potential outlier point in the regression model; when removed, the model and interaction term remained significant (*p* = 0.02 and *p* = 0.01, respectively; not shown). **E.** Expression of dopamine receptor 2 (*Drd2*) explained 39.1% (R^2^) of the variation in response latency after the requirement shift in the attentional set-shifting task (*p* = 0.02). The slope of the regression lines were significantly different between A1221 males and Veh controls. **F.** Expression of dopamine transport gene *Slc6a3* explained 24.6% (R^2^) of the variation in response latency after the requirement shift in the attentional set-shifting task (*p* = 0.05). The slopes of the regression lines were significantly different between A1221 males and Veh controls.

#### Estradiol and Dopamine cells

In the subset of females used for IHC, serum E2 was predictive of the number of dopamine cells (DA; i.e., mean TH+ cells) in the VTA in a manner that interacted with Treatment group (Figure 6C). The overall model explained 38.7% of the variation in the data, with an adjusted R^2^ of 27.3% (*F*(3,16) = 3.39, *p* = 0.04). The relationship between E2 and VTA DA was negative (β = -0.56, STE = 0.25, *t* = -2.28, *p* = 0.03); E2 interacted positively with Treatment and VTA DA (β = 0.97, STE = 0.42, *t* = 2.31, *p* = 0.03), indicating that E2 was a positive predictor of VTA DA in Veh controls, but not in A1221-exposed females. This relationship was unique to VTA in females: in the SN, the overall regression model between E2 and DA was not significant (*p* > 0.1), and E2 did not predict DA cell numbers in male rats (*p* > 0.1 for both VTA and SN).

#### Estradiol and gene expression

There were no significant predictive relationships in females or males between E2 and expression of the target genes (all models *p* > 0.1).

#### Dopamine cells and behavior

There were no significant predictive relationships in females or males between VTA and SN DA cells and behavioral outcomes (all models *p* > 0.1).

#### Gene expression and behavior

Of the target genes, *Drd1, Drd2, and Slc6a3* varied with behavioral outcomes in a way that differed between treatment groups only in male rats (all other gene-behavior models *p* > 0.1). *Drd1* predicted preference for sucrose solution between male treatment groups (Figure 6D): the overall model explained 48.8% of the variation in the data, with an adjusted R^2^ of 39.1% (*F*(3,16) = 5.08, *p* = 0.01). *Drd1* interacted positively with Treatment and sucrose preference (β = 13.88, STE = 4.50, *t* = 3.09, *p* = 0.007), indicating *Drd1* increased with sucrose preference differently between A1221-exposed males and Veh controls. Notably, the effect of *Drd1* on sucrose preference was not driven by a single data point: when a potential outlier was removed, the model (*F*(3,15) = 4.47, *p* = 0.02; adjusted R^2^ = 36.6%) and interaction effect (β = 7.78, STE = 2.65, *t* = 2.94, *p* = 0.01) remained significant. *Drd2* and *Slc6a3* both predicted latency to respond in the attentional set-shifting task between male treatment groups (Figure 6E,F). For *Drd2*, the overall model explained 44.6% of the variation in the data, with an adjusted R^2^ of 34.3% (*F*(3,16) = 4.30, *p* = 0.02). *Drd2* interacted positively with Treatment and response latency (β = 0.69, STE = 0.28, *t* = 2.58, *p* = 0.02), indicating that *Drd2* increased with response latency in Veh controls but not A1221-exposed males. Similarly for *Slc6a3*, the overall model explained 36.5% of the variation in the data, with an adjusted R^2^ of 24.6% (*F*(3,16) = 3.07, *p* = 0.05). *Slc6a3* interacted positively with Treatment and response latency (β = 0.61, STE = 0.29, *t* = 2.04, *p* = 0.05), indicating that *Slc6a3* increased with response latency in Veh controls but not A1221-exposed males.

## Discussion

The presented data demonstrate that perinatal exposure to the PCB mixture A1221 has sex-specific effects on DA-mediated behavior and gene expression in DA-producing regions of the midbrain. Moreover, it provides evidence that different mechanisms drive these responses in male and female rats. Female rats exposed to A1221 perinatally showed broader behavioral effects including a depressive-like behavioral phenotype accompanied with decreased response latency in the set-shifting task. Males had limited behavioral effects – reduced response latency – but also higher expression of the estrogen receptor alpha gene *Esr1* and non-significant effects on DA signaling genes in the midbrain. In females, E2 was a modest predictor of behavioral responses and DAergic cells in the midbrain. In males, expression of multiple DAergic genes in the midbrain predicted behavioral responses. Taken together, these results suggest that behavioral disruptions associated with perinatal A1221 exposure rely partially on hormonal mechanisms in female rats and on neuromolecular mechanisms in male rats.

### Depressive-like behavior, dopamine, and estradiol

Anhedonia, or the reduced ability to experience pleasure, is a key aspect of major depression disorder (First, 2013); reductions in preference for sucrose solution is considered a proxy of anhedonic behavior in rodents (Primo et al., 2023). Sucrose preference and anhedonia are modulated in part by DAergic activity and the DA-producing regions of the midbrain: in rodents, DA depletion in the SN or in the VTA reduce sucrose preference (Martínez-Hernández et al., 2006; Santiago et al., 2014; Shibata et al., 2009). DA receptors 1 and 2 (D1 and D2, respectively) also impact sucrose preference: antagonists of either receptor decrease preference for lower concentrations of sucrose (Muscat & Willner, 1989), and D1-deficient mice show reduced motivation for sucrose reward (El-Ghundi et al., 2003). E2 has known antidepressive effects in rodents (Carrier et al., 2015; Romano-Torres & Fernández-Guasti, 2010). When E2 availability is reduced via ovariectomy, female rodents show decreased preference for sucrose (Cheng et al., 2013; Jie, 2023). Estrogen receptors modulate DA synthesis and are found in DAergic cells of the midbrain (Creutz & Kritzer, 2002), which may partially explain E2’s effects on appetitive and affective behaviors.

We used the weakly estrogenic PCB mixture A1221, which we have previously shown increases serum E2 levels in developing but not adult female rats (Streifer et al., 2023), and increases *Drd1* in the hypothalamus of male rats (Liberman et al., 2020). To our knowledge, this is the first evidence that perinatal exposure to A1221 decreases sucrose preference in female rats – a depressive-like phenotype – and that E2 influences DAergic cell numbers in the VTA of these animals differently after A1221 exposure. A similar relationship was observed between E2 and performance on the set-shifting task, suggesting that female rats exposed to A1221 may utilize E2 differently than controls. Interestingly, we did not observe any differences in expression of estrogen receptors or DA signaling genes in female rats, nor any relationships between gene expression and behavior. E2 exerts tonic inhibition on D2 receptors to stimulate sucrose intake (Bâ et al., 2018); it is possible that A1221 exposure changes the way E2 interacts with DAergic targets in the midbrain, thereby impacting functionality of these cells and affecting behavior without changing the overall measurement of these targets. In males, we did not see rote effects of A1221 on sucrose preference; however, the positive relationship between *Drd1* and sucrose preference in A1221 males may suggest that A1221 increases functional activity of DA receptor 1 – which is associated with higher sucrose preference (Muscat & Willner, 1989).

### Response latency and dopamine

PCB mixtures reduce DA transporter levels, synaptosomal DA content, and induce DAergic cell death in adult exposure models (Bemis & Seegal, 2004; Caudle et al., 2006; D. W. Lee et al., 2012). In the current study, both female and male rats exposed perinatally to A1221 exhibited reduced response latency in the set-shifting task. Male A1221 rats had higher expression of *Esr1* in the midbrain DA producing regions, and A1221 exposure non-significantly impacted expression of *Drd1* and *Slc6a3*. Of these, DA transporter genes are targets of PCBs in adult exposure models and both *Drd1* and *Slc6a3* have known roles in affective and attentional processing (Andersen & Teicher, 2000; El-Ghundi et al., 2003; Krause, 2008; Richardson & Miller, 2004). *Drd2* and *Slc6a3* had negative relationships with response latency in the A1221 males; higher D2 is associated with lower response inhibition in humans and rodents (Beste et al., 2016; Eagle et al., 2011; Robertson et al., 2015), and antagonism of this receptor increases response latency in the attentional set-shifting task (Haluk & Floresco, 2009).

Increased response latency is considered a measure of cognitive decline in aging rodents and may be reflective of higher cognitive effort in complex tasks (Young et al., 2010), but the inverse of this – decreased response latency – is more difficult to interpret. It is possible that A1221 improves performance in the set-shifting task; however, the reduced response latency without concurrent improvement in cognitive flexibility (i.e., ability to shift response requirements) is more suggestive of a hyperactive or disinhibited response type (Bari & Robbins, 2013; Puumala et al., 1996). D2 receptor and dopamine transporter genes are implicated in the etiology of ADHD (DiMaio et al., 2003; Swanson et al., 2000; Volkow et al., 2009), and D2 specifically is implicated in hyperactivity (Fan et al., 2010). The relationship observed here – higher *Drd2* and *Slc6a3* associated with decreased response latency in A1221 males – provides some evidence for ADHD-like symptomology in these animals. However, identifying whether this represents a hyperactive phenotype or one of behavioral disinhibition requires more nuanced testing using appropriate paradigms, such as the 5-choice serial reaction time or Go/No-Go tests (Loos et al., 2010).

### A note on conditioned orienting

A core inspiration for this research was to determine if disrupted hormonal processes in early developmental life may impact DAergic behaviors and neuro-molecular function. Of the selected behaviors, conditioned orienting is a DA-mediated behavior with a sex-specific presentation that does not rely on availability of E2 in adults (Hilz et al., 2021, 2022; H. J. Lee et al., 2011). We hypothesized that sex differences in conditioned orienting may be organized in early life, and that exposure to A1221 in this period may impact the development of the orienting phenotype. Our results did not support this hypothesis. In separate analyses not shown here, we examined the relationships between OR phenotype (i.e., OR level at the end of conditioning, classified as Orienters or Nonorienters using the methods from Hilz, Lewis, et al., 2019) and various behavioral outcomes. We found strong associations across multiple endpoints, including a positive correlation with DA cell numbers in the midbrain and impaired performance in attentional set-shifting tasks. However, the experiment was not powered to consider both OR phenotype and exposure to A1221 in concert. The results taken as they are suggest that A1221 in the perinatal window did not target processes involved in the development of the orienting phenotype – and the limited cognitive-attentional effects in general suggest that perinatal exposure to A1221 may not be a candidate for the development of ADHD-like behavior in rodents. Other exposure models, such as inter- and transgenerational models, or the use of chemical mixtures that have different impacts on hormonal processes like anti-androgenic compounds or complex EDC mixtures (Gore et al., 2022), may help elucidate the mechanisms that organize attentional phenotypes.

## Conclusion

Taken together, the current data provide new evidence that exposure to A1221 during the perinatal period affects developmental processes that influence behavioral function in adulthood, and that the mechanisms via which this occurs may differ between male and female rats. These results are consistent with previous work from our lab on EDCs reporting that they 1) have broader behavioral effects in female rats compared to males (Gillette et al., 2022; Gore et al., 2022); 2) cause hormonally mediated behavioral disruptions in females (Reilly et al., 2015); and 3) involve neuromolecular-mediated disruptions, more so in male rats (Gore et al., 2021, 2022). To fully understand the implications of EDC exposure on attentional function and the increasing comorbidities between neuropsychiatric disorders, it is crucial to assess impacts in models that approximate real-world scenarios. Understanding the mechanisms that drive behavioral effects of EDCs can help design better interventions in a world with an increasing burden of EDC exposure.

